# High-Resolution Linkage Map and QTL Analyses for Machine Harvest Traits in Autotetraploid Blueberry

**DOI:** 10.1101/2020.05.13.093633

**Authors:** Francesco Cappai, Rodrigo R. Amadeu, Juliana Benevenuto, Ryan Cullen, Alexandria Garcia, Adina Grossman, Luís Felipe V. Ferrão, Patricio Munoz

## Abstract

Blueberry (*Vaccinium corymbosum* and hybrids) is an autotetraploid crop whose commercial relevance has been growing steadily during the last twenty years. However, the ever-increasing cost of labor for hand-picking blueberry is one main constraint in competitive marketing of the fruit. Machine harvestability is, therefore, a key trait for the blueberry industry. Understanding the genetic architecture of traits through quantitative trait locus (QTL) mapping is the first step towards implementation of molecular breeding for faster genetic gains. Despite recent advances in software development for autotetraploid genetic mapping, a high-resolution map is still not available for blueberry. In this study, we crafted a map for autotetraploid low-chill highbush blueberry containing 11,292 SNP markers and a total size of 1,953.97 cM (average density of 5.78 markers/cM). This map was subsequently used to perform QTL analyses for traits relevant to machine harvesting: firmness, firmness retention, and fruit detachment force. Significant QTL peaks were identified for all the traits. The QTL intervals were further explored for putative candidate genes. Genes related to cell wall remodeling were highlighted in the firmness and firmness retention intervals. For fruit detachment force, transcription factors involved in fruit abscission were detected. Altogether, our findings provide the basis for future fine-mapping and molecular breeding efforts for machine harvesting in blueberry.

## 1 INTRODUCTION

It is not uncommon to find plants with different ploidy levels within the same genus. In fact, such chromosomal alterations are not only tolerated by plant genomes, but can also drive evolution, for example, promoting speciation (Adams and Wendel, 2005). Polyploid organisms are classified as either allopolyploids or autopolyploids, depending on the degree of divergence between their subgenomes (Brubaker et al., 1999). Autopolyploids, such as blueberry (*Vaccinium corymbosum* L.), potato (*Solanum tuberosum* L.), and alfalfa (*Medicago sativa* L.), contain multiple copies of the same chromosome set, which can all pair and exchange genetic material during gamete formation. In contrast, allopolyploids, such as wheat (*Triticum aestivum* L.), coffee (*Coffea arabica* L.), and strawberry (*Fragaria ananassa* Duch.) contain two or more divergent genomes, which usually show bivalent pairing and have disomic inheritance, like diploid organisms. While in diploid systems the study of allelic inheritance is relatively simple, polysomic inheritance in autopolyploids increases the number of possible genetic configurations and impacts downstream genetic analyses, including linkage map construction and quantitative trait loci (QTL) mapping (Bever and Felber, 1992).

QTL mapping is the association study between phenotypic and genetic variants usually performed by tracking recombination events of the possible QTL along chromosomes (Broman and Sen, 2009). In breeding programs, QTL analysis has been historically used to understand the genetic basis of complex traits, which can ultimately support the implementation of marker-assisted selection. To this end, the development of a reliable linkage map and haplotype inference are both considered crucial steps. A classical approach for autopolyploids relies on building individual maps at each homolog level using a combination of diploid software and single-dose markers, i.e. makers that segregate in a diploid fashion (1:1 and 1:2:1) (Mollinari and Garcia, 2019). Although widely adopted this strategy has a few shortcomings, including the use of only a subset of markers, the low-power to detect markers in repulsion, and the use *ad-hoc* procedures to assemble the Linkage Groups (LGs) into homology groups. Just recently, new methods addressing autotetraploid genetics and higher orders of segregation patterns have been reported. Software such TetraploidSNPmap (Hackett et al., 2017), an updated version of the TetraploidMap (Hackett et al., 2007), polymapR (Bourke et al., 2018), and mappoly ((Mollinari and Garcia, 2019) overcame those limitations and allowed the construction of high-density linkage maps using SNP data and multi-dose markers. Potato (Da Silva et al., 2017), sweetpotato (*Ipomoea batatas* L.) (Mollinari et al., 2020) and forage grasses (Ferreira et al., 2019; Deo et al., 2020) are examples of autopolyploid crops that have benefited from the utilization of such modern methodologies in QTL inference and linkage mapping analyses. However, to the best of our knowledge, no high-density linkage map has been built for autotetraploid blueberries. Linkage maps reported for blueberry have relied on diploid populations (Rowland et al., 2014) and/or used a small number of markers and individuals (McCallum et al., 2016; Schlautman et al., 2018). A high-density linkage map has only been built for cranberry (*Vaccinium macrocarpon* Ait.) (Schlautman et al., 2017), a diploid-relative of blueberry.

Highbush blueberry is an autotetraploid crop (2n = 4x = 48), considered the second most important soft fruit, after strawberry (Brazelton et al., 2017). Despite the considerable dedicated world acreage and varietal improvement through traditional breeding, blueberry still remains one of the most expensive commonly sold fruits by weight (USDA Economic Research Service, 2020). A main factor contributing to these sustained prices is the high production cost, mostly due to the laborious hand-picking process, which can represent more than 50% of the total production cost (Ehlenfeldt and Martin, 2002; Olmstead and Finn, 2014; Gallardo et al., 2018). In addition, a number of countries, such as the USA, are experiencing labor shortages, further exacerbating this issue (Roka and Guan, 2018). A possible solution that has been tested in blueberry fields is the mechanization of the harvesting operations. Mechanical harvesting of berries can lead to a considerable (up to 80%) reduction in cost, while also reducing food-borne illnesses caused by improper produce handling by laborers (Berger et al., 2010; Olmstead and Finn, 2014). Machine harvesting consists of metal rods shaking the blueberry bushes, forcing berries to fall. To successfully implement machine harvest, the following traits, alongside with plant architecture, are crucial: I) high berry firmness to withstand physical damage; II) high detachability in ripe (blue) fruit; and III) low detachability in unripe (green) fruit.

Firmness is a genetically controlled trait that has long been among the focus of blueberry breeding programs (Edwards et al., 1974; Cappai et al., 2018; Cellon et al., 2018; Ferrão et al., 2018). It is also a desirable trait across the whole market chain: for growers, less fruit is discarded due to physical damage; for retailers, firmer berries have a longer shelf life; and for consumers, firm texture positively correlates to liking (Mehra et al., 2013; Olmstead and Finn, 2014; Yu et al., 2014; Gilbert et al., 2015). At a molecular level, fruit firmness is intertwined with fruit ripening, and it is the result of a complicated network of interactions between two main hormones, ethylene and abscisic acid (ABA), and the cell wall disassembly process (Brummell, 2006; Vicente et al., 2007; Chiabrando and Giacalone, 2011; Cappai et al., 2018). The exact cascade of events that leads to fruit softening has not been completely elucidated in blueberry, even though current evidence indicates that pectin solubilization in the cell wall could be one of the main outcomes (Lara et al., 2004; Vicente et al., 2007; Angeletti et al., 2010; Beaudry et al., 2016). Fruit Detachment Force (FDF), or abscission force, is the force required for a single berry to detach from a stem at the point of the pedicel junction. Machine harvesting requires a large differential between the FDF of green and ripe berries in order to maximize the number of ripe berries that detach from the plant at a defined force point (Malladi et al., 2012; Vashisth et al., 2015). In addition, the force required to detach ripe berries should also be low in absolute terms to ensure efficient fruit removal while avoiding excessive damage to the plant itself. Molecular mechanisms underlying fruit abscission have been extensively studied in tomato (*Solanum lycopersicum* L.) and *Arabidopsis*, which pointed to the role of MADS-box family transcription factors on the regulation of the pedicel-abscission zone development (Ferrándiz, 2002; Nakano et al., 2012; Ito and Nakano, 2015). Cell wall-modifying proteins and programmed cell death at the abscission zone have been shown to play a role in the cell separation processes (Roberts et al., 2002; Tsuchiya et al., 2015). There is also strong evidence for the interplay between phytohormones in regulating fruit abscission, with ethylene and auxin enhancing and inhibiting the process, respectively (Estornell et al., 2013; Patterson et al., 2016). In blueberry, comparison of two genotypes contrasting in FDF levels provided preliminary insights into differential expression of genes related to phytohormone pathways (Vashisth et al., 2015).

Here, we aimed to develop a high-resolution linkage map for autotetraploid blueberry and perform QTL mapping for machine harvest-related traits to reveal their genetic architecture. To this end, we phenotyped a large mapping population for berry firmness and FDF, and genotyped using a high marker density. Genomic regions associated with the phenotypes were further explored for functional insights. Our results are promising for marker-assisted selection implementation and future fine-mapping to identify causal variants and elucidate the molecular basis underlying these traits.

## 2 MATERIALS AND METHODS

### 2.1 Plant growth conditions

The population used in this study consisted of 237 individuals planted in 2010 derived from a cross between two varieties, ‘Sweetcrisp’ and ‘Indigocrisp’, both released by the Blueberry Breeding Program at the University of Florida (Lyrene, 2009, 2016). All plants were grown in Waldo, Florida (29°24’30.2”N 82°08’32.6”W) at a spacing of approximately 86 cm within rows and 2 m between rows. They received 5.68 L of water per plant a day injected with 180 ppm sulfuric acid through a drip irrigation system to adjust soil pH. During the growing season (February through October), a 10-5-5 liquid fertilizer was also applied through the irrigation system. Plants were pruned in the summer months of June and July. Insecticide for spotted wing drosophila as well as fungicides (Pristine, Switch, and Proline) were applied six times a year to manage diseases and crop damage. In winter months (December and January), freeze protection measures were applied as necessary.

### 2.2 Plant genotyping

Genomic DNA was extracted in 2018 from leaf tissue of each individual in the mapping population and both parental cultivars using a 2 % CTAB extraction method (Xin and Chen, 2012). Library preparation and sequencing were performed by RAPiD Genomics (Gainesville, FL, USA) using a sequence capture approach. Briefly, 6,000 custom-designed biotinylated probes of 120-mer were selected based on the distribution on the ‘Draper’ reference genome (Benevenuto et al., 2019), and previously tested on the parental cultivars. Sequencing of the entire population was carried out in the Illumina HiSeq platform using 150 cycle paired-end runs. Raw reads were demultiplexed and barcodes were removed. Data was cleaned and trimmed at the 3’ end by removing bases with quality scores lower than 20 and reads with more than 10% of the bases with quality scores lower than 20, using Fastx Toolkit v.0.0.14 http://hannonlab.cshl.edu/fastx_toolkit/. Trimmed reads were aligned to the ‘Draper’ blueberry reference genome using MOSAIK v. 2.2.30 (Lee et al., 2014). The 12 largest chromosomes of each homeologous set from the ‘Draper’ genome were used as references (Colle et al., 2019). SNPs were called using FreeBayes v.1.0.1 (Garrison and Marth, 2012), targeting the 6,000 probe regions. SNPs were further filtered by I) minimum mapping quality of 30; II) mean depth of coverage of 40; III) maximum missing data of 50%; IV) minor allele frequency of 0.01; V) only biallelic loci; and VI) no monomorphism. A total of 21,513 SNPs were kept, after these filtering steps. Subsequently, read counts per allele and individual were extracted from the variant call file using vcftools v.0.1.16 (Danecek et al., 2011). The updog R package was used to call the tetraploid allele dosages based on the read counts and ‘f1’ model (Gerard et al., 2018). The “f1” model implemented in the software uses parental information as a baseline for estimating the allele dosages of the progenies. The posterior probability means per SNP for each individual were rounded toward the closest integer and used as our genotypes.

### 2.3 Plant phenotyping

#### 2.3.1 Firmness

Mature fruits were collected at peak ripeness, in March and early April of 2018 and 2019. Berries were stored at 4°C overnight, brought to room temperature, and their firmness was measured using the FirmtechTM 2 firmness tester (BioWorks, Wamego, KS). Berries were placed on the machine with their equatorial axis perpendicular to the instrument’s surface and probe. The force-deformation response of compressed berries was measured in grams per 1 mm of deflection (g/mm) (Ehlenfeldt and Martin, 2002). A sample size of 25 berries per genotype was used in 2018 and a sample size of 50 berries per genotype was used in 2019, as this sample size was reported to increase accuracy (Cappai et al., 2018). Berry firmness measurements were averaged per genotype within each year.

#### 2.3.2 Firmness retention

In 2019, berries previously used to measure firmness were stored for four additional weeks in a climate-controlled chamber at 4°C in the dark. They were then removed from the chamber, brought to room temperature, and firmness was measured as described above. Firmness retention was computed as the deviation between firmness computed after one day of storage and firmness computed after four week of storage.

#### 2.3.3 Fruit detachment force and differential fruit detachment force

Fruit detachment force (FDF) was measured in 2019 for ripe blueberries and unripe green berries using a DS2 Digital Force Gauge (Imada, Northbrook, Ill.) (Olmstead and Finn, 2014). Berries were placed in the prong of the force gauge and pulled at a 90° angle away from the bush. At the time of berry separation at the pedicel-berry junction, the force measurement was recorded in Newtons. If a berry fell from the bush without being pulled, a force measurement of 0.1 N was recorded. The sample size for fruit detachment force was 25 green berries and 25 ripe berries for each parent and offspring in the mapping population. The differential fruit detachment force (or ΔFDF) was calculated per genotype as follows: ΔFDF= (average FDF of ripe berries) – (average FDF of green berries).

### 2.4 Phenotypic analysis

We computed adjusted means (LS-means) for each genotype based on a linear model, where genotype and year were considered fixed effects; LS-means for each trait were used as our response variables for the subsequent QTL mapping analyses. To compute genomic heritabilities, we used linear mixed models to estimate the variance components using restricted maximum likelihood estimator approach in the asreml-R software (Butler, 2019). For the firmness trait, we fit the following linear mixed model *y* = *Xb* + *Zg* + *ɛ*, where *y* is the response variable; *b* is the fixed effect of year; *g* is the random effect for the genotype nested within the year where *g* ~ *MV N* (0, *G*_*b*_ ⊗ *K*_*g*_), being *G*_*b*_ a 2 × 2 unstructured variance-covariance matrix for the year effect and *K*_*g*_ the genomic relationship matrix; *ɛ* is the residual effects where *ɛ* ~ *MV N* (0, Σ_*b*_ ⊗ *I*_*g*_), being Σ_*b*_ a 2 × 2 unstructured variance-covariance matrix for the residual term associated with the year variation, and *I*_*g*_ an identity matrix. *X* and *Z* are the fixed and random effects, respectively. *K*_*g*_ was estimated by the average of the genomic relationship matrices computed with the identity-by-descent probabilities for every 1 cM with the genotypic probabilities derived from the linkage map. Heritability (*h*^2^) was computed within each year as 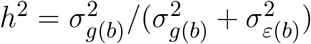. For FDF, ΔFDF, and firmness retention, *h*^2^ was computed based on a similar model, but with no year effect.

### 2.5 Map crafting and parameters

The linkage map was built based on the method proposed by (Mollinari and Garcia, 2019) using the mappoly software and following a series of filtering steps as follows. After genotyping, markers in small unassembled contigs were removed (21,513 remaining markers). Marker genotypes with a probability of correct dosage assignment lower than 0.80 were treated as missing data. Markers with higher than 20% of missing information across individuals were also removed (14,820 remaining markers). Individuals with more than 10% of marker information missing were removed (237 remaining individuals). Finally, markers with Mendelian segregation distortion were removed, considering a chi-square test with Bonferroni correction assuming an alpha level of significance of 0.05 (14,792 remaining markers). Pairwise recombination fractions for all the possible linkage phases between pairs of markers were estimated. The phasing configuration with the highest logarithm of the odds (LOD) score was used to build a matrix of recombination fractions. Based on the recombination fraction heatmap, we performed clustering analyses with the unweighted pair group method with arithmetic mean algorithm to assign 12 clusters for our marker data. Each cluster would represent one of the 12 blueberry linkage groups (LG). The marker group assignment clusters was compared with the genome mapping results (i.e. on which chromosome the marker mapped, based on the ‘Draper’ genome assembly). Mismatching markers were removed, resulting in 14,598 markers, which were grouped into 12 distinct LGs. Within each LG, the markers were ordered using the ‘Draper’ reference genome. With the recombination fraction heatmap, the overall map order was checked using the multidimensional scaling algorithm as proposed by Preedy and Hackett (2016), and was also visually inspected (Margarido et al., 2007). Both methodologies showed similar results. Therefore, we kept the mapping, since the multidimensional scaling order is not powerful enough to solve local inversions (shorter range distance) and would introduce ordering errors (Preedy and Hackett, 2016; Mollinari et al., 2020).

Multipoint recombination fraction and haplotype phasing were estimated using Hidden Markov Models (HMM) (Mollinari and Garcia, 2019). Briefly, the HMM procedure starts the chain with the first 20 markers, estimates all the possible phases, selects the one with the highest likelihood, then, in order, adds one marker at a time, and reevaluates the map likelihood and distance. We chose a combination of parameters for the HMM procedure so that when a new marker is included, it minimizes the chance of inflating the map while keeping the chain open for other possible phasing configurations and maintaining a high phasing accuracy in a feasible computation time. In the mappoly software, we set the following HMM parameters: `start.set = 20, thres.twopt = 10, thres.hmm = 10, extend.tail = 200, info.tail = TRUE, sub.map.size.diff.limit = 10, phase.number.limit = 20, reestimate.single.ph.confirguration = TRUE, tol = 10e-3`, and `tol.final = 10e-4`. If the recombination fraction heatmap or the multidimensional scaling graphics showed possible inversions and/or misplacements of a map segment, alternative orders (chromosome rearrangements) were also evaluated with the HMM procedure and finally, the order with the highest likelihood was kept as the final order. Markers that inflated the map were manually removed during the process. Map density was evaluated graphically with LinkageMapView (Ouellette et al., 2018).

### 2.6 QTL mapping

The QTL mapping was performed with a random effect interval mapping using a simplified approach derived from (Pereira et al., 2020). First, within the final linkage map, we computed the conditional probabilities of each individual haplotype in relation to the 36 possible haplotype combinations of an autotetraploid biparental cross for every 1 cM of the map. To this end, we used an HMM procedure adapted from (Lander and Green, 1987) and estimated in the mappoly software (Mollinari and Garcia, 2019). The QTL mapping was performed for each trait using these haplotype probabilities as our predictor variables in a random-effect interval mapping adapted for autopolyploids. Considering *n* individuals, the random effects model for one QTL can be written as *y* = *μ* + *g*_*q*_ + *ɛ*, where *y* is the response variable, *μ* is the overall mean effect (fixed), *g*_*q*_ is a vector *n* × 1 with the individual random effects for the locus *q*, where 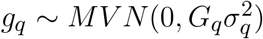 being *G*_*q*_ the genetic relationship matrix for this given locus, where *G*_*q*_ = *Z*_*q*_Π*Z*′_*q*_, *Z*_*q*_ is an incidence matrix *n* × 36 containing the genotype conditional probabilities of the locus *q*, and Π is a 36 × 36 matrix containing the expected proportion of shared alleles by identity-by-descent between every genotype pair, 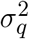 is the QTL variance in the locus *q*, and 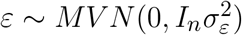. After fitting a given model for a given QTL, the linear score statistic was computed, which provided a p-value for the given QTL that was then converted to its logarithm as *LOP* = −*log*_10_(*p* – *value*]. To determine the significance threshold for each trait, we performed 1,000 rounds of permutations. In each round, the phenotypic observations were shuffled, a random-effect interval mapping model was performed, and the highest LOP values were extracted. Based on the LOP values, we computed the 95% quantile of the second and third peak across all permutations (Doerge and Churchill, 1996; Chen and Storey, 2006). For comparison, we also performed the QTL mapping using fixed-effects interval mapping as proposed by Hackett et al. (2001). The QTL heritability was computed based on the variance estimates as 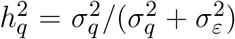. The individuals best linear unbiased predictions (BLUPs) were decomposed in order to estimate each allelic effect and the combination of alleles for each parent. The qtlpoly software was used to perform QTL analysis (Pereira et al., 2020). Support intervals for the QTL location were computed based on LOP 1-drop rule, similar to the standard LOD 1-drop rule (Lander and Botstein, 1989; Li, 2011).

### 2.7 Candidate gene screening

Intervals previously selected for each QTL peak based on the LOP 1-drop rule were explored in order to find candidate genes (Lander and Botstein, 1989; Li, 2011). For firmness, measured over the course of 2 years, overlapping QTL intervals were merged and the whole resulting area was further analyzed.

Predicted genes at the QTL intervals in the ‘Draper’ genome were extracted and the correspondent predicted protein sequences were retrieved (Colle et al., 2019). Functional annotation was performed using BLASTp (v.2.9.0) search against the eudicots non-redundant database with an e-value cut-off of 10^−5^ (Altschul et al., 1990). Domain and gene ontology terms were annotated using InterProScan v 5.35-74.0 (Quevillon et al., 2005).

## 3 RESULTS

We genotyped 237 individuals and obtained 21,513 SNPs markers. After filtering, we were able to create a tetraploid linkage map that included 11,292 SNP markers with a total size of 1,953.97 cM. This means an average density of 5.78 markers/cM (Table 1). Markers were well distributed throughout the 12 linkage groups (Figure 1 and Supplementary Figure 1). We found only two gaps between adjacent mapped markers higher than 15 cM (17.01 cM on LG 6 and 16.89 cM on LG 9). The length of each LG ranged from 128.45 cM to 193.78 cM, with an average of 162.83 cM. During map construction, multidimensional scaling graphical results showed two possible order mismatches between our mapping population and the ‘Draper’ genome. The order mismatches occurred at LG 2 and LG 9 in distal chromosomic segments comparing the reference genome with the suggested multidimensional scaling order (Supplementary Figure 1). All possible orders and inversions of the individual segments were tested, and the one with the highest likelihood was set as the true order for the map. LGs with their marker order, positions in cM, and parental phasing can be found in Supplementary Data 1.

**Table 1.** Summary statistics of the blueberry map showing the linkage groups (LG), total number of markers assembled (Markers), LG length in cM, log-likelihood of the LG (LL), and total number of gaps assuming three different thresholds (5, 10, and 15 cM).

| LG | Markers | Length | LL | Total gaps |  |  |
| --- | --- | --- | --- | --- | --- | --- |
|  |  |  |  | >5 cM | >10 cM | >15 cM |
| 1 | 932 | 150.40 | -9,360.05 | 1 | 0 | 0 |
| 2 | 1,093 | 189.12 | -10,724.5 | 3 | 1 | 0 |
| 3 | 1,060 | 168.28 | -10,551.79 | 3 | 0 | 0 |
| 4 | 826 | 153.23 | -8,961.25 | 2 | 0 | 0 |
| 5 | 1,129 | 187.46 | -11,084.64 | 3 | 2 | 0 |
| 6 | 795 | 154.01 | -8,094.59 | 4 | 1 | 1 |
| 7 | 749 | 128.45 | -7,231.92 | 2 | 0 | 0 |
| 8 | 911 | 149.05 | -10,013.13 | 1 | 0 | 0 |
| 9 | 889 | 193.78 | -9,367.19 | 3 | 1 | 1 |
| 10 | 955 | 151.38 | -9,569.56 | 2 | 0 | 0 |
| 11 | 1,085 | 175.09 | -11,101.22 | 0 | 0 | 0 |
| 12 | 868 | 153.72 | -9,638.22 | 1 | 0 | 0 |
| Total | 11,292 | 1,953.97 | -115,698.06 | 25 | 5 | 2 |

**Figure 1.**
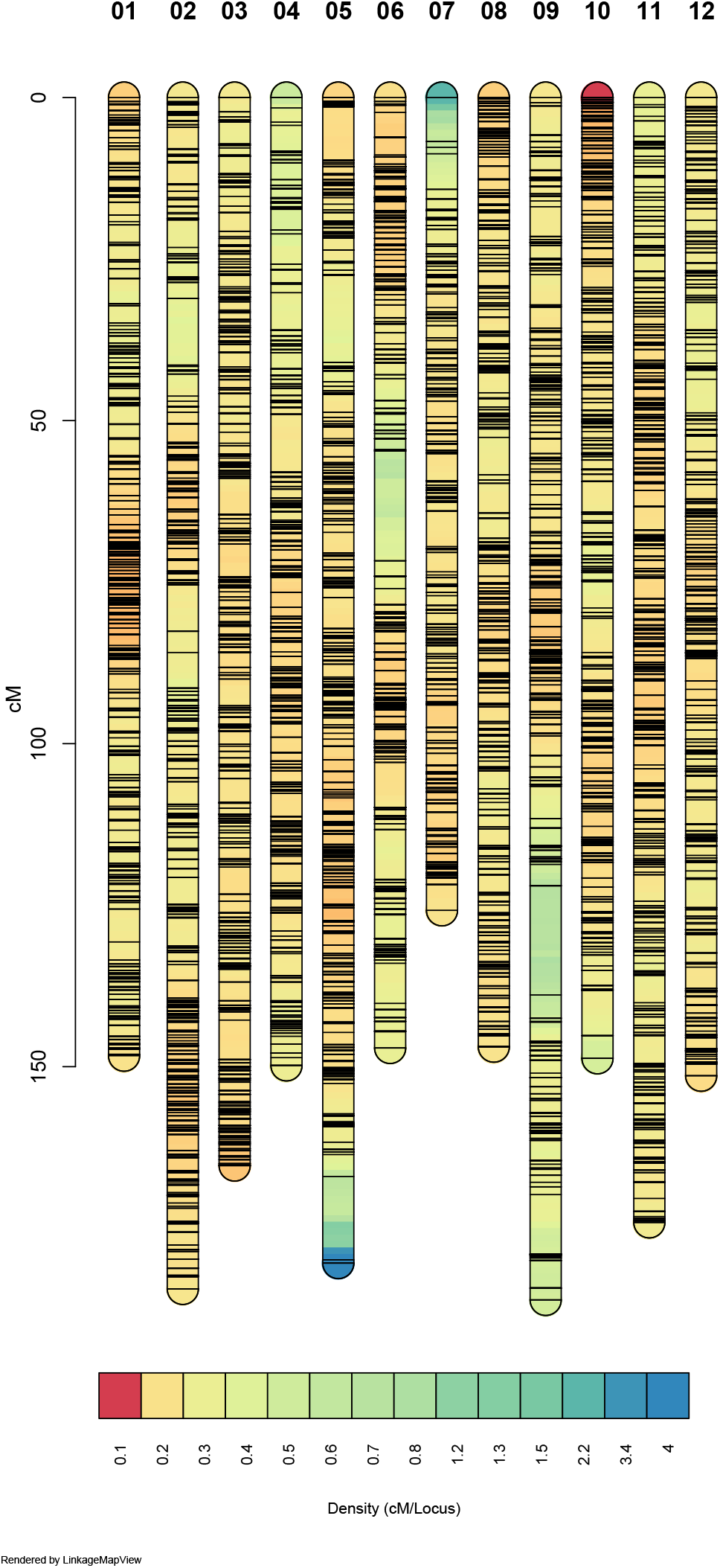
Linkage map density. The 12 linkage groups are represented and colored according to the marker density. Warmer colors represent denser marker region and each black line shows the marker position. Figure built with LinkageMapView software.

Variance estimation for each trait was obtained using linear mixed models. Heritability ranged from 0.34 for FDF to 0.75 for firmness retention traits (Table 2). For the QTL mapping, the average threshold computed via permutations for the second and third highest peaks had a LOP score of 3.12, and 2.71 respectively. In total, we mapped five QTLs spanning three different LGs in the blueberry genome. Notably, for all traits, we observed individual QTLs explaining more than 15% of the phenotypic variation. For firmness retention, a single QTL on LG 8 explained 27% of the phenotypic variance. For firmness, the location of the QTL reported was consistent across both years (Figure (2 and Table 3). The QTL mapping LOP profile was similar to the fixed-effects interval mapping model (Supplementary Figure 2).

**Table 2.** Phenotypic averages 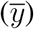 for the parents ‘Sweetcrisp’, ‘Indigocrisp’, and their offspring, estimated genotypic 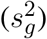 and error 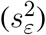 variances, and genotypic heritability (*h*^2^) for each trait. NA when not measured.

| Trait | $\bar{y}_{sweet}$ | $\bar{y}_{indigo}$ | $\bar{y}_{offspring}$ | $s_g^2$ | $s_e^2$ | $h^2$ |
| --- | --- | --- | --- | --- | --- | --- |
| Firmness 2018 | 259.71 | 238.06 | 237.7 | 298.04 | 356.76 | 0.46 |
| Firmness 2019 | 283.26 | 267.46 | 271.52 | 569.29 | 549.37 | 0.51 |
| Firmness retention | NA | NA | 8.62 | 563.38 | 192.14 | 0.75 |
| Ripe FDF | 0.43 | 0.74 | 0.84 | 0.02 | 0.04 | 0.34 |
| $\Delta$ FDF | -0.87 | -1.34 | 1.93 | 0.11 | 0.22 | 0.34 |

**Figure 2.**
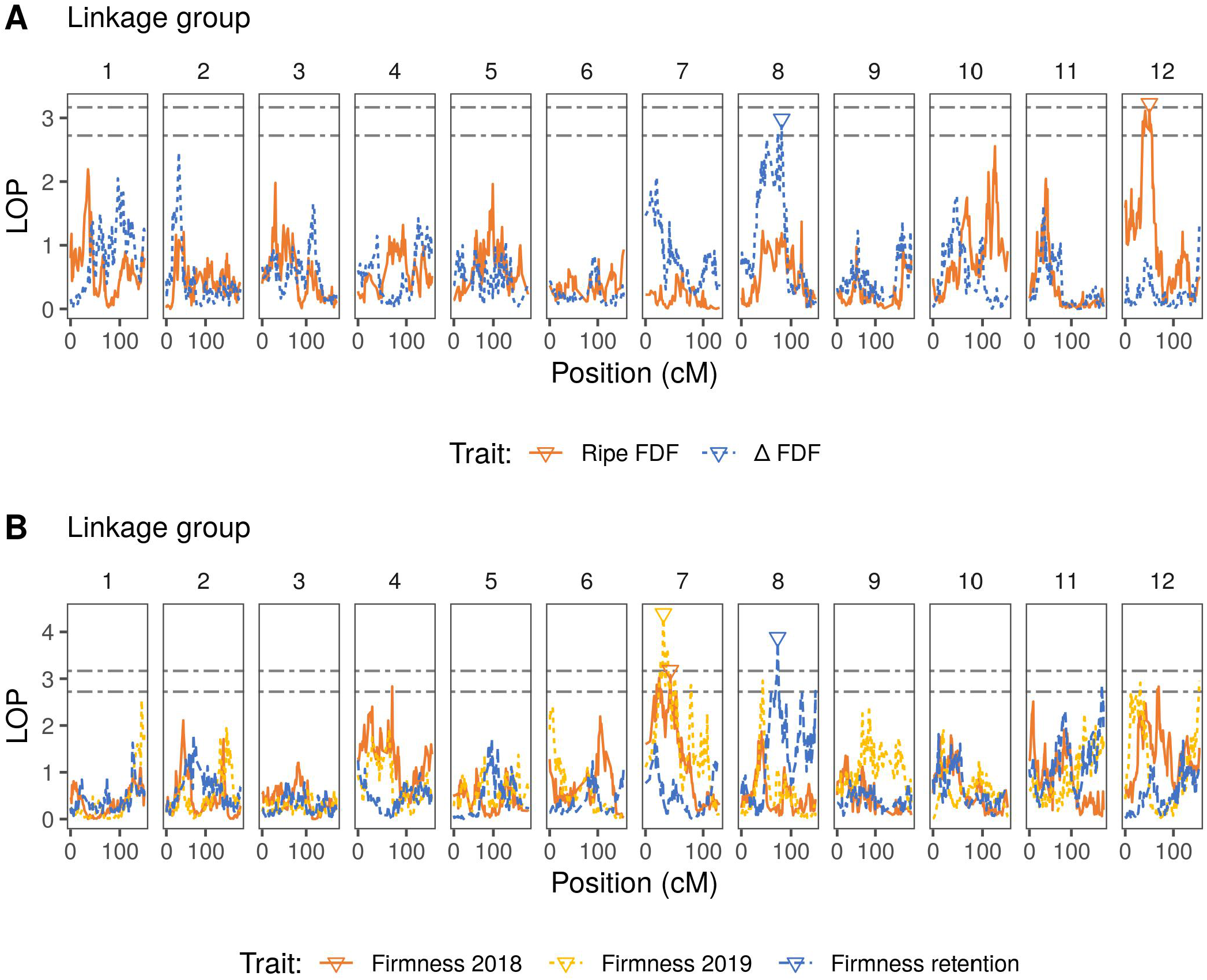
QTL mapping profiles for blueberry machine harvesting traits throughout the 12 linkage groups. LOP is the negative logarithm score of the p-values for the presence of the QTL in the model. Triangles indicate inferred QTL positions for each trait. Black dashed lines represent the permutation thresholds considering 95% quantile of the second (top) and third (bottom) highest peak. **(A)** Fruit detachment force profiles for ripe fruit and for the difference (Δ) between ripe and unripe fruit. **(B)** Firmness after one day of storage (for 2018 and 2019 seasons) and firmness retention (difference between 2019 firmness after one day of storage and after four weeks of storage).

**Table 3.** QTL position inferred for each trait, linkage group (LG), highest logarithm of the p-value of the QTL (LOP), the peak position in the map in centiMorgans (Peak) with confidence interval assuming 1-LOP drop rule (Lower and Upper bound), SNP position with the maximum LOP score of the peak (Loci), and the QTL heritability (*h*^2^).

| Trait | LG | LOP | Peak (cM) | Lower (cM) | Upper (cM) | Loci | $h^2$ |
| --- | --- | --- | --- | --- | --- | --- | --- |
| Firmness 2018 | 7 | 3.08 | 43.13 | 15.06 | 53.39 | 12_11134097 | 0.16 |
| Firmness 2019 | 7 | 4.29 | 31.11 | 26.42 | 38.38 | 12_9191799 | 0.16 |
| Firmness retention | 8 | 3.78 | 73.01 | 69.51 | 75.47 | 13_14221092 | 0.27 |
| Ripe FDF | 12 | 3.13 | 50.04 | 36.44 | 55.79 | 22_10452951 | 0.19 |
| $\Delta$ FDF | 8 | 2.88 | 81.26 | 39.11 | 83.16 | 13_16002128 | 0.16 |

The biological significance of predicted genes identified within the QTL intervals was investigated considering *in silico* functional annotation (Supplementary Data 2). For firmness and firmness retention, we highlighted genes involved in calcium binding, transport and signaling pathways (e.g. *calcium-transporting ATPase 1*, *calcium uniporter protein 4*, *CBL-interacting serine/threonine-protein kinase 8*, *calcium-binding protein CML41*), and cell-wall modifying enzymes (e.g. *endo-1,4-beta-xylanase*, *beta-galactosidase 3*, *mannan endo-1,4-beta-mannosidase 7*, *galacturonosyltransferase-like 1*, *UDP-glucuronate:xylan alpha-glucuronosyltransferase 2*, *leucine-rich repeat extensin-like protein*). In the QTL interval for FDF of ripe berries and ΔFDF, we detected genes related to MADS-box transcription factors involved in fruit abscission, such as *FUL-like protein/AGL8* and *AGL15*. Since ripening is a fundamental process for all traits in this study, genes involved in hormone-induced senescence could also be plausible candidates. Several genes involved in phytohormone pathways and transport were detected, e.g. for ethylene (*ethylene responsive transcription factor ERF011* and *ERF114-like*), abscisic acid (*abscisic acid 8’-hydroxylase 4*, *molybdenum cofactor sulfurase ABA3*, *protein C2-DOMAIN ABA-RELATED 4*), auxin (s*erine/threonine-protein kinase D6PKL2*, *ARF guanine-nucleotide exchange factor GNOM*, *indole-3-pyruvate monooxygenase YUCCA6*), among others (Supplementary Data 2).

## 4 DISCUSSION

Blueberry production costs are subjected to increasing high hand-harvesting prices. Machine harvesting is an economically viable solution that requires plants with particular characteristics, such as firm berries and fruits that easily detach only when ripe (Olmstead and Finn, 2014). Despite their importance, the genetic basis underlying the variation of these machine harvest-related traits remain poorly understood in blueberry. Our contributions in this paper are three-fold: I) we crafted a high-density genetic map for low chill (also known as southern highbush) autotetraploid blueberry; II) we mapped potential QTLs associated with machine harvest-related traits, showing their location and effect; and III) we provided some insights into potential molecular mechanisms for firmness and FDF based on predicted gene functions at significant QTL intervals.

To date, most of the blueberry genetic maps reported in the literature have relied on diploid methods, ignoring the fact that allele dosages, polysomic segregation, and multilocus information should be considered in the analyses for a more robust inference. By combining such information with the current genomic resources available, we have created a unifying framework to develop high-quality genetic maps that could be used, for example, as a scaffolding strategy to accomplish future genome assembly projects. Furthermore, the map provides a statement about the inheritance pattern involved in the transmission of genes aiding in the task of haplotype inference. Although other genetic maps were reported in blueberry, high-density polyploid maps are still lacking, which have been already reported for other autopolyploid crop species (Da Silva et al., 2017; Ferreira et al., 2019; Schlautman et al., 2018; Deo et al., 2020; Mollinari et al., 2020). Moreover, previous blueberry linkage maps were not assembled considering prior genomic information was based on smaller populations, lower marker density and/or, more importantly, did not consider most of the genetic particularities that are intrinsic to a tetraploid system (Rowland et al., 2014; McCallum et al., 2016; Schlautman et al., 2018). While our map shows map inflation, this is lower than has been found in another genetic map built with similar methodology (Mollinari et al., 2020). Three main reasons can be pointed out for this apparent inflation: I) misplacement of the marker order, II) genotyping errors, and III) genotyping dosage errors (Mollinari and Garcia, 2019). Aside from differences in species ploidy level, the genotyping quality and the methodology used for allele dosage calling are also potential reasons to explain the difference in map inflation between these two species.

In this report, we present the first high-density map for tetraploid blueberry (*Vaccinium* spp), which brings new advances for the blueberry community but also opens questions that warrant further investigation. First, we detected order mismatches between our map and the previously released ‘Draper’ genome (Colle et al., 2019) in LGs 2 and 9. These could be due to real chromosomal translocations specific to these genotypes, genome assembling issues, or mapping artifacts. Another possible investigation is regarding double-reduction landscape in blueberry. Even though there is no strong evidence of its presence (Krebs and Hancock, 1989; Amadeu et al., 2016; McCallum et al., 2016), with such high marker density, double-reduction phenomena could also be investigated extensively in blueberry as it is being done in potato (Hackett et al., 2013; Bourke et al., 2015).

After the genetic map construction, QTL mapping was performed to understand the genetic architecture of machine harvest-related traits. By estimating the number, position, and effect of markers underling phenotypic variation, QTL results open new opportunities for marker-assisted selection implementation, which can ultimately accelerate and maximize genetic gains. Despite their potential contribution to breeding programs, phenotype-genotype association studies in blueberry have not been fully utilized. Only recently were the first genome-wide association studies (GWAS) reported for fruit quality and aroma traits (Ferrão et al., 2018, 2020), while QTL mapping has been performed only for chilling requirement and cold hardiness traits (Rowland et al., 2014). Herein, using a random-effect model for multiple-QTL mapping, we identified four major QTLs associated with four important traits related to machine harvesting in blueberry. Markers associated with QTLs explained low-to-moderate portions of the phenotypic variances, which suggest a quantitative nature of these traits. In accordance to this, previous GWAS analyses in blueberry also detected few and scattered associations between SNPs and fruit firmness using dominant gene action models, which also explained a small proportion of the trait variation (Ferrão et al., 2018; Benevenuto et al., 2019).

QTL regions using the LOP 1-drop rule spanned genomic intervals from 5.96 to 44.05 cM, encompassing from 1.68 to 9.71 Mb in the reference genome, with each interval harboring hundreds of genes. Given the large size of linkage blocks inherent in most QTL mapping populations, it is difficult at this stage to point out specific candidate genes underlying the variability of the traits. However, a number of biologically plausible candidate genes can be highlighted. Overall, all traits considered in this study are tightly related to fruit ripening, senescence, and abscission. These processes are the result of complex interactions of plant hormones, signaling pathways, and transcriptional and cellular modifications (Seymour et al., 2002; Giovannoni, 2004; Estornell et al., 2013; Cappai et al., 2018).

For firmness and firmness retention traits, the most striking candidates found at the QTL intervals were those related to cell wall modification, such as glycosyltransferases (EC 2.4), glycosylases (EC 3.2) and leucine-rich repeat extensin-like proteins. Fruit softening, at both *in planta* and in storage conditions, has been mainly associated with depolymerization and solubilization of hemicellulose and pectin in the cell wall (Huber, 1983; Brummell, 2006; Chen et al., 2015).

For ripe FDF and ΔFDF traits, genes potentially related to the transcriptional regulation of the differentiation and activation of the abscission zone are interesting candidates, such as *FRUITFULL/AGL8* and *AGL15*. *FRUITFULL* (*FUL*) is a MADS-BOX transcription factor associated with the differentiation of the dehiscence zone of the silique in *Arabidopsis* (Ferrándiz et al., 2000), and also has significant sequence similarities with the tomato gene *MACROCALYX* (Nakano et al., 2012). The constitutive expression of *FRUITFULL* has been shown to be sufficient to prevent formation of the dehiscence zone (Ferrándiz et al., 2000). *AGAMOUS like 15* (*AGL15*) is also a MADS-BOX transcription factor that maintains a non-senescent state of plant tissues, and whose constitutive expression resulted in delayed floral organ abscission in *Arabidopsis* (Fernandez et al., 2000; Estornell et al., 2013). Other transcription factors detected at the ΔFDF QTL region could also have an indirect role in abscission, such as *OFP7* (Wang et al., 2013), *SPATULA* (Heisler et al., 2001; Girin et al., 2011), *TCP13* (Koyama et al., 2007), and *HHO5* (Butenko and Simon, 2015; Moreau et al., 2016).

Several genes involved in phytohormone pathways were detected at the QTL intervals for all traits. For firmness and firmness retention traits, ethylene and ABA are well-known for their role as ripening promoters and, subsequently, senescence and softening (Suzuki et al., 1997; Zhu et al., 2012; Sun et al., 2013; Cappai et al., 2018). Within the QTL interval for firmness, for example, there were two putative ‘ethylene-responsive transcription factor (ERFs)’ encoding genes. A fine QTL mapping in tomato also detected an ERF gene underlying firmness variation and its increased expression was associated with soft fruit texture in the tomato mapping population (Chapman et al., 2012). Genes related to phytohormone pathways could also be involved in fruit abscission and, therefore, underlying FDF and ΔFDF traits (Lipe and Morgan, 1972; Riov et al., 1990; Chauvaux et al., 1997; Estornell et al., 2013). Despite the copious literature supporting the role of hormones in fruit softening and dehiscence, their pathways and effects are extremely complex and interconnected with other hormones, so many candidates can be speculated to have a role. In addition to hormone-related genes, a diverse set of calcium-related genes were also found in the QTL regions, including some with potential binding, transport, and calcium-activated signal transduction functions. Calcium also has well-documented roles in signaling, water relations, and cell wall modification during fruit ripening in various fruit crops; therefore, these genes may also be underlying the variation of the traits (Conway and Sams, 1987; García et al., 1996; Pilar Hernandez et al., 2006; Ciccarese et al., 2013; Beaudry et al., 2016; Munir et al., 2016; Gao et al., 2019).

Altogether, we reported a high-quality linkage map and candidate QTL regions for four machine harvest-related traits in autotetraploid blueberry. We used state-of-art algorithms for the linkage analysis applied to polyploids and, therefore, our results can be relevant for the polyploid community. We detected QTL associated with machine harvesting traits and characterized their genetic architecture and the potential for marker-assisted selection implementation in the breeding program. Finally, we highlighted some plausible candidate genes at the QTL intervals. Future efforts to identify causal genes and variants can include a combination of fine mapping, transcriptomics, and functional testing of the genes, such as CRISPR-Cas9 inactivation.

## 5 NOMENCLATURE

FDF: Fruit Detachment Force
LG: Linkage Group
GWAS: Genome-Wide Association Study
HMMs: Hidden Markov Models
QTL: Quantitative Trait Loci
SNP: Single Nucleotide Polymorphism
LOD: Logarithm of the Odds
LOP: Logarithm of the p-value
ABA: Abscisic Acid

## AUTHOR CONTRIBUTIONS

PM and FC conceived and designed the study. FC, RC, AGA, and AGR conducted the field experiment, phenotyping, and DNA extraction. RRA and LFVR performed statistical genetics analyses. FC and JB performed the SNP filtering and gene annotation. FC and RRA wrote the manuscript. All authors read, reviewed and approved the final version of this manuscript.

## ACKNOWLEDGMENTS

We thank Dr. Marcelo Mollinari and Dr. Guilherme da Silva Pereira for their valuable comments during mapping and QTL analyses. We also thank Lauren Scott and Werner Collante for their support and guidance in the laboratory.

## SUPPLEMENTAL DATA

For this preprint version is available upon request to the authors.

**Supplementary Data 1.** It contains 11 columns by 11,292 rows, where rows are the markers and columns present the assembled linkage group (LG), SNP name (scaffold + position), position in cM, parent 1 phase (a, b, c, d homologues), parent 2 phase (e, f, g, h). ‘ ’ represents the alternative SNP allele, and ‘o’ otherwise (reference allele).

**Supplementary Data 2.** It contains four spreadsheets, one per trait, where rows are the predicted genes in QTL intervals, and columns have in silico annotation information: transcript ID, blastp description, protein length, e-value, mean percentage of identity, GO terms, GO description, enzyme codes, enzyme names, InterPro domains, and manual annotations, respectively.

## REFERENCES

Adams, K. L. and Wendel, J. F. (2005). Polyploidy and genome evolution in plants. Curr. Opin. Plant Biol. 8, 135–141. doi:10.1016/J.PBI.2005.01.001

Altschul, S. F., Gish, W., Miller, W., Myers, E. W., and Lipman, D. J. (1990). Basic local alignment search tool. J Mol. Bio. 215, 403–410. doi:10.1016/S0022-2836(05)80360-2

Amadeu, R. R., Cellon, C., Olmstead, J. W., Garcia, A. A. F., Resende, M. F. R., and Muñz, P. R. (2016). AGHmatrix: R package to construct relationship matrices for autotetraploid and diploid species: A blueberry example. The Plant Genome 9, 1–10. doi:10.3835/plantgenome2016.01.0009

Angeletti, P., Castagnasso, H., Miceli, E., Terminiello, L., Concellón, A., Chaves, A., et al. (2010). Effect of preharvest calcium applications on postharvest quality, softening and cell wall degradation of two blueberry (*Vaccinium corymbosum*) varieties. Postharvest Biol. Technol. 58, 98–103. doi:10.1016/j.postharvbio.2010.05.015

Beaudry, R. M., Hanson, E. J., Beggs, J. L., and Beaudry, R. M. (2016). Applying calcium chloride postharvest to improve highbush blueberry firmness. HortScience 28, 2–4. doi:10.21273/HORTSCI.28.10.1033

Benevenuto, J., Ferrão, V. L. F., Amadeu, R. R., and Munoz, P. (2019). How can a high-quality genome assembly help plant breeders? GigaScience 8. doi:10.1093/gigascience/giz068

Berger, C. N., Sodha, V. S., Shaw, R. K., Griffin, P. M., Pink, D., Hand, P., et al. (2010). Fresh fruit and vegetables as vehicles for the transmission of human pathogens. Environ. Microbiol. 12, 2385–2397. doi:10.1111/j.1462-2920.2010.02297.x

Bever, J. D. and Felber, F. (1992). The theoretical population genetics of autopolyploidy. In Oxford surveys in evolutionary biology volume 8, eds. D. Futuyma and J. Antonovics (Oxford University Press). 185–217

Bourke, P. M., van Geest, G., Voorrips, R. E., Jansen, J., Kranenburg, T., Shahin, A., et al. (2018). polymapR — linkage analysis and genetic map construction from F1 populations of outcrossing polyploids. Bioinformatics 34, 3496–3502. doi:10.1093/bioinformatics/bty371

Bourke, P. M., Voorrips, R. E., Visser, R. G. F., and Maliepaard, C. (2015). The double-reduction landscape in tetraploid potato as revealed by a high-density linkage map. Genetics 201, 853–863. doi:10.1534/genetics.115.181008

Brazelton, C., Kayla, Y., and Bauer, N. (2017). Global Blueberry Statistics and Intelligence Report. Tech. rep., International Blueberry Organization

Broman, K. W. and Sen, S. (2009). A Guide to QTL Mapping with R/qtl (Springer)

Brubaker, C. L., Paterson, A. H., and Wendel, J. F. (1999). Comparative genetic mapping of allotetraploid cotton and its diploid progenitors. Genome 42, 184–203

Brummell, D. A. (2006). Cell wall disassembly in ripening fruit. Funct. Plant Biol. 33, 103–119. doi:10.1071/FP05234

Butenko, M. A. and Simon, R. (2015). Beyond the meristems: similarities in the CLAVATA3 and INFLORESCENCE DEFICIENT IN ABSCISSION peptide mediated signalling pathways. J. Exp. Bot. 66, 5195–5203. doi:10.1093/jxb/erv310

Butler, D. (2019). asreml: fits the linear mixed model. R Package (VSN International)

Cappai, F., Benevenuto, J., Ferrão, L., and Munoz, P. (2018). Molecular and genetic bases of fruit firmness variation in blueberry — a review. Agronomy 8, 174. doi:10.3390/agronomy8090174

Cellon, C., Amadeu, R. R., Olmstead, J. W., Mattia, M. R., Ferrao, L. F. V., and Munoz, P. R. (2018). Estimation of genetic parameters and prediction of breeding values in an autotetraploid blueberry breeding population with extensive pedigree data. Euphytica 214, 87. doi:10.1007/s10681-018-2165-8

Chapman, N. H., Bonnet, J., Grivet, L., Lynn, J., Graham, N., Smith, R., et al. (2012). High-resolution mapping of a fruit firmness-related quantitative trait locus in tomato reveals epistatic interactions associated with a complex combinatorial locus. Plant Physiol. 159, 1644–1657. doi:10.1104/pp.112.200634

Chauvaux, N., Child, R., John, K., Ulvskov, P., Borkhardt, B., Prinsen, E., et al. (1997). The role of auxin in cell separation in the dehiscence zone of oilseed rape pods. J. Exp. Bot. 48, 1423–1429. doi:10.1093/jxb/48.7.1423

Chen, H., Cao, S., Fang, X., Mu, H., Yang, H., Wang, X., et al. (2015). Changes in fruit firmness, cell wall composition and cell wall degrading enzymes in postharvest blueberries during storage. Scientia Horticulturae 188, 44–48. doi:10.1016/j.scienta.2015.03.018

Chen, L. and Storey, J. D. (2006). Relaxed significance criteria for linkage analysis. Genetics 173, 2371–2381. doi:10.1534/genetics.105.052506

Chiabrando, V. and Giacalone, G. (2011). Shelf-life extension of highbush blueberry using 1-methylcyclopropene stored under air and controlled atmosphere. Food Chem. 126, 1812–1816. doi:10.1016/j.foodchem.2010.12.032

Ciccarese, A., Stellacci, A. M., Gentilesco, G., and Rubino, P. (2013). Effectiveness of pre- and post-veraison calcium applications to control decay and maintain table grape fruit quality during storage. Postharvest Biol. Technol. 75, 135–141. doi:10.1016/j.postharvbio.2012.08.010

Colle, M., Leisner, C. P., Wai, C. M., Ou, S., Bird, K. A., Wang, J., et al. (2019). Haplotype-phased genome and evolution of phytonutrient pathways of tetraploid blueberry. GigaScience 8. doi:10.1093/gigascience/giz012

Conway, W. S. and Sams, C. E. (1987). The effects of postharvest infiltration of calcium, magnesium, or strontium on decay, firmness, respiration, and ethylene production in apples. J. Amer. Soc. Hort. Sci.

Da Silva, W. L., Ingram, J., Hackett, C. A., Coombs, J. J., Douches, D., Bryan, G. J., et al. (2017). Mapping loci that control tuber and foliar symptoms caused by PVY in autotetraploid potato (*Solanum tuberosum* l.). G3 (Bethesda) 7. doi:10.1534/g3.117.300264

Danecek, P., Auton, A., Abecasis, G., Albers, C. A., Banks, E., DePristo, M. A., et al. (2011). The variant call format and VCFtools. Bioinformatics 27, 2156–2158. doi:10.1093/bioinformatics/btr330

Deo, T. G., Ferreira, R. C. U., Lara, L. A. C., Moraes, A. C. L., Alves-Pereira, A., de Oliveira, F. A., et al. (2020). High-resolution linkage map with allele dosage allows the identification of regions governing complex traits and apospory in guinea grass (*Megathyrsus maximus*). Front. Plant Sci. 11. doi:10.3389/fpls.2020.00015

Doerge, R. W. and Churchill, G. A. (1996). Permutation tests for multiple loci affecting a quantitative character. Genetics 142, 285–294

Edwards, T. J., Sherman, W., and Sharpe, R. (1974). Evaluation and inheritance of fruit color, size, scar, firmness and plant vigor in blueberry. HortScience 9, 20–22

Ehlenfeldt, M. K. and Martin, R. B. (2002). A survey of fruit firmness in highbush blueberry and species-introgressed blueberry cultivars. HortScience 37, 386–389. doi:10.21273/HORTSCI.37.2.386

Estornell, L. H., Agustí, J., Merelo, P., Talón, M., and Tadeo, F. R. (2013). Elucidating mechanisms underlying organ abscission. Plant Sci. 199, 48–60. doi:10.1016/j.plantsci.2012.10.008

Fernandez, D. E., Heck, G. R., Perry, S. E., Patterson, S. E., Bleecker, A. B., and Fang, S.-C. (2000). The embryo MADS domain factor AGL15 acts postembryonically: inhibition of perianth senescence and abscission via constitutive expression. Plant Cell 12, 183–197. doi:10.1105/tpc.12.2.183

Ferreira, R. C. U., Lara, L. A. d. C., Chiari, L., Barrios, S. C. L., do Valle, C. B., Valério, J. R., et al. (2019). Genetic mapping with allele dosage information in tetraploid *Urochloa decumbens* (Stapf) R. D. Webster reveals insights into spittlebug (*Notozulia entreriana* Berg) resistance. Front. Plant Sci. 10. doi:10.3389/fpls.2019.00092

Ferrándiz, C. (2002). Regulation of fruit dehiscence in *Arabidopsis*. J. Exp. Bot. 53, 2031–2038

Ferrándiz, C., Liljegren, S. J., and Yanofsky, M. F. (2000). Negative regulation of the SHATTERPROOF genes by FRUITFULL during *Arabidopsis* fruit development. Science 289, 436–438. doi:10.1126/science.289.5478.436

Ferrão, L. F. V., Johnson, T. S., Benevenuto, J., Edger, P. P., Colquhoun, T. A., and Munoz, P. R. (2020). Genome-wide association of volatiles reveals candidate loci for blueberry flavor. New Phytol. Early View. doi:10.1111/nph.16459

Ferrão, V. L. F., Benevenuto, J., Oliveira, I. d. B., Cellon, C., Olmstead, J., Kirst, M., et al. (2018). Insights into the genetic basis of blueberry fruit-related traits using diploid and polyploid models in a GWAS context. Front. Ecol. Evol. 6, 107. doi:10.3389/fevo.2018.00107

Gallardo, R. K., Stafne, E. T., DeVetter, L. W., Zhang, Q., Li, C., Takeda, F., et al. (2018). Blueberry producers’ attitudes toward harvest mechanization for fresh market. HortTechnology 28, 10–16. doi:10.21273/HORTTECH03872-17

Gao, Q., Xiong, T., Li, X., Chen, W., and Zhu, X. (2019). Calcium and calcium sensors in fruit development and ripening. Scientia Horticulturae 253, 412–421. doi:10.1016/j.scienta.2019.04.069

García, J. M., Herrera, S., and Morilla, A. (1996). Effects of postharvest dips in calcium chloride on strawberry. J. Agric. Food Chem. 44, 30–33. doi:10.1021/jf950334l

Garrison, E. and Marth, G. (2012). Haplotype-based variant detection from short-read sequencing. arXiv doi:arXiv:1207.3907

Gerard, D., Ferrão, L. F. V., Garcia, A. A. F., and Stephens, M. (2018). Genotyping polyploids from messy sequencing data. Genetics 210, 789–807. doi:10.1534/GENETICS.118.301468

Gilbert, J. L., Guthart, M. J., Gezan, S. A., De Carvalho, M. P., Schwieterman, M. L., Colquhoun, T. A., et al. (2015). Identifying breeding priorities for blueberry flavor using biochemical, sensory, and genotype by environment analyses. PLoS ONE 10, e0138494. doi:10.1371/journal.pone.0138494

Giovannoni, J. J. (2004). Genetic regulation of fruit development and ripening. Plant Cell 16, S170–S180. doi:10.1105/tpc.019158

Girin, T., Paicu, T., Stephenson, P., Fuentes, S., Körner, E., O’Brien, M., et al. (2011). INDEHISCENT and SPATULA interact to specify carpel and valve margin tissue and thus promote seed dispersal in *Arabidopsis*. Plant Cell 23, 3641–3653. doi:10.1105/tpc.111.090944

Hackett, C. A., Boskamp, B., Vogogias, A., Preedy, K. F., and Milne, I. (2017). TetraploidSNPMap: Software for linkage analysis and QTL mapping in autotetraploid populations using SNP dosage data. Heredity 108, 438–442. doi:10.1093/jhered/esx022

Hackett, C. A., Bradshaw, J. E., and McNicol, J. W. (2001). Interval mapping of quantitative trait loci in autotetraploid species. Genetics 159, 1819–1832

Hackett, C. A., McLean, K., and Bryan, G. J. (2013). Linkage analysis and QTL mapping using SNP dosage data in a tetraploid potato mapping population. PLoS ONE 8, e63939. doi:10.1371/journal.pone.0063939

Hackett, C. A., Milne, I., Bradshaw, J. E., and Luo, Z. (2007). TetraploidMap for Windows: Linkage map construction and QTL mapping in autotetraploid species. Heredity 98, 727–729. doi:10.1093/jhered/esm086

Heisler, M. G., Atkinson, A., Bylstra, Y. H., Walsh, R., and Smyth, D. R. (2001). SPATULA, a gene that controls development of carpel margin tissues in *Arabidopsis*, encodes a bHLH protein. Development 128, 1089–1098

Huber, D. J. (1983). Polyuronide degradation and hemicellulose modifications in ripening tomato fruit. J. Amer. Soc. Hort. Sci. 108, 405–409

Ito, Y. and Nakano, T. (2015). Development and regulation of pedicel abscission in tomato. Front. Plant Sci. 6, 442. doi:10.3389/fpls.2015.00442

Koyama, T., Furutani, M., Tasaka, M., and Ohme-Takagi, M. (2007). TCP transcription factors control the morphology of shoot lateral organs via negative regulation of the expression of boundary-specific genes in *Arabidopsis*. Plant Cell 19, 473–484. doi:10.1105/tpc.106.044792

Krebs, S. L. and Hancock, J. F. (1989). Tetrasomic inheritance of isoenzyme markers in the highbush blueberry, *Vaccinium corymbosum* L. Heredity 63, 11–18. doi:10.1038/hdy.1989.70

Lander, E. S. and Botstein, S. (1989). Mapping mendelian factors underlying quantitative traits using RFLP linkage maps. Genetics 121, 185

Lander, E. S. and Green, P. (1987). Construction of multilocus genetic linkage maps in humans. Proc. Natl. Acad. Sci. U. S. A. 84, 2363–2367. doi:10.1073/pnas.84.8.2363

Lara, I., García, P., and Vendrell, M. (2004). Modifications in cell wall composition after cold storage of calcium-treated strawberry (*Fragaria × ananassa* Duch.) fruit. Postharvest Biol. Technol. 34, 331–339. doi:10.1016/j.postharvbio.2004.05.018

Lee, W.-P., Stromberg, M. P., Ward, A., Stewart, C., Garrison, E. P., and Marth, G. T. (2014). MOSAIK: a hash-based algorithm for accurate next-generation sequencing short-read mapping. PLoS ONE 9. doi:10.1371/journal.pone.0090581

Li, H. (2011). A quick method to calculate QTL confidence interval. J. Genetics 90, 355–360

Lipe, J. A. and Morgan, P. W. (1972). Ethylene: role in fruit abscission and dehiscence processes. Plant Physiol. 50, 759–764. doi:10.1104/pp.50.6.759

Lyrene, P. M. (2009). ‘Sweetcrisp’ southern highbush blueberry plant (USA: USPP20027P3)

Lyrene, P. M. (2016). Blueberry plant named ‘FL98-325’ (USA: US20150106980P1)

Malladi, A., Vashisth, T., and Johnson, L. K. (2012). Ethephon and methyl jasmonate affect fruit detachment in rabbiteye and southern highbush blueberry. HortScience 47, 1745–1749. doi:10.21273/HORTSCI.47.12.1745

Margarido, G. R. A., Souza, A. P., and Garcia, A. A. F. (2007). OneMap: software for genetic mapping in outcrossing species. Hereditas 144, 78–79. doi:10.1111/j.2007.0018-0661.02000.x

McCallum, S., Graham, J., Jorgensen, L., Rowland, L. J., Bassil, V. N., Hancock, J. F., et al. (2016). Construction of a SNP and SSR linkage map in autotetraploid blueberry using genotyping by sequencing. Mol. Breed. 36, 41. doi:10.1007/s11032-016-0443-5

Mehra, L. K., MacLean, D. D., Savelle, A. T., and Scherm, H. (2013). Postharvest disease development on southern highbush blueberry fruit in relation to berry flesh type and harvest method. Plant Dis. 97, 213–221. doi:10.1094/PDIS-03-12-0307-RE

Mollinari, M. and Garcia, A. A. F. (2019). Linkage analysis and haplotype phasing in experimental autopolyploid populations with high ploidy level using hidden Markov models. G3 (Bethesda) 9, 3297–3314. doi:10.1534/g3.119.400378

Mollinari, M., Olukolu, B. A., Da Pereira, G. S., Khan, A., Gemenet, D., Craig Yencho, G., et al. (2020). Unraveling the hexaploid sweetpotato inheritance using ultra-dense multilocus mapping. G3 (Bethesda) 10, 281–292. doi:10.1534/g3.119.400620

Moreau, F., Thévenon, E., Blanvillain, R., Lopez-Vidriero, I., Franco-Zorrilla, J. M., Dumas, R., et al. (2016). The Myb-domain protein ULTRAPETALA1 INTERACTING FACTOR 1 controls floral meristem activities in *Arabidopsis*. Development 143, 1108–1119. doi:10.1242/dev.127365

Munir, S., Khan, M. R. G., Song, J., Munir, S., Zhang, Y., Ye, Z., et al. (2016). Genome-wide identification, characterization and expression analysis of calmodulin-like (CML) proteins in tomato (*Solanum lycopersicum*). Plant Phys. Biochem. 102, 167–179. doi:10.1016/j.plaphy.2016.02.020

Nakano, T., Kimbara, J., Fujisawa, M., Kitagawa, M., Ihashi, N., Maeda, H., et al. (2012). MACROCALYX and JOINTLESS interact in the transcriptional regulation of tomato fruit abscission zone development. Plant Physiol. 158, 439–450. doi:10.1104/pp.111.183731

Olmstead, J. W. and Finn, C. E. (2014). Breeding highbush blueberry cultivars adapted to machine harvest for the fresh market. HortTechnology 24, 290–294. doi:10.21273/HORTTECH.24.3.290

Ouellette, L. A., Reid, R. W., Blanchard, S. G., and Brouwer, C. R. (2018). LinkageMapView-rendering high-resolution linkage and QTL maps. Bioinformatics 34, 306–307. doi:10.1093/bioinformatics/btx576

Patterson, S. E., Bolivar-Medina, J. L., Falbel, T. G., Hedtcke, J. L., Nevarez-McBride, D., Maule, A. F., et al. (2016). Are we on the right track: can our understanding of abscission in model systems promote or derail making improvements in less studied crops? Front. Plant Sci. 6, 1268. doi:10.3389/fpls.2015.01268

Pereira, G. S., Gemenet, D. C., Mollinari, M., Olukolu, B. A., Wood, J. C., Mosquera, V., et al. (2020). Multiple QTL mapping in autopolyploids: a random-effect model approach with application in a hexaploid sweetpotato full-sib population. Genetics doi:10.1534/genetics.120.303080

Pilar Hernandez, M., Maria, J., and Rafael, G. (2006). Effect of calcium dips and Chitosan coating on postharvest life of strawberry (*Fragaria × ananasa*). Postharvest Biol. Technol. 39, 247–253. doi:10.1016/j.postharvbio.2005.11.006

Preedy, K. F. and Hackett, C. A. (2016). A rapid marker ordering approach for high-density genetic linkage maps in experimental autotetraploid populations using multidimensional scaling. Theor. Appl. Genet. 129, 2117–2132. doi:10.1007/s00122-016-2761-8

Quevillon, E., Silventoinen, V., Pillai, S., Harte, N., Mulder, N., Apweiler, R., et al. (2005). InterProScan: protein domains identifier. Nucleic Acids Res. 33, W116–W120. doi:10.1093/nar/gki442

Riov, J., Dagan, E., Goren, R., and Yang, S. F. (1990). Characterization of abscisic acid-induced ethylene production in citrus leaf and tomato fruit tissues. Plant Physiol. 92, 48–53. doi:doi.org/10.1104/pp.92.1.48

Roberts, J. A., Elliott, K. A., and Gonzalez-Carranza, Z. H. (2002). Abscission, dehiscence, and other cell separation processes. Annu. Rev. Plant Biol. 53, 131–158. doi:10.1146/annurev.arplant.53.092701.180236

Roka, F. M. and Guan, Z. (2018). Farm labor management trends in Florida, USA-challenges and opportunities. Intl. J. Agric. Manage. 7, 1–9. doi:10.22004/ag.econ.292479

Rowland, L. J., Ogden, E. L., Bassil, N., Buck, E. J., McCallum, S., Graham, J., et al. (2014). Construction of a genetic linkage map of an interspecific diploid blueberry population and identification of QTL for chilling requirement and cold hardiness. Mol. Breed. 34, 2033–2048. doi:10.1007/s11032-014-0161-9

Schlautman, B., Covarrubias-Pazaran, G., Diaz-Garcia, L., Iorizzo, M., Polashock, J., Grygleski, E., et al. (2017). Construction of a high-density american cranberry (*Vaccinium macrocarpon* Ait.) composite map using genotyping-by-sequencing for multi-pedigree linkage mapping. G3 (Bethesda) 7, 1177–1189. doi:10.1534/g3.116.037556

Schlautman, B., Diaz-Garcia, L., Covarrubias-Pazaran, G., Schlautman, N., Vorsa, N., Polashock, J., et al. (2018). Comparative genetic mapping reveals synteny and collinearity between the American cranberry and diploid blueberry genomes. Mol. Breed. 38. doi:10.1007/s11032-017-0765-y

Seymour, G. B., Manning, K., Eriksson, E. M., Popovich, A. H., and King, G. J. (2002). Genetic identification and genomic organization of factors affecting fruit texture. J. Exp. Bot. 53, 2065–2071. doi:10.1093/jxb/erf087

Sun, Y., Hou, Z., Su, S., and Yuan, J. (2013). Effects of ABA, GA3 and NAA on fruit development and anthocyanin accumulation in blueberry. J. South China Agr. Univ. 34, 6–11

Suzuki, A., Kikuchi, T., and Aoba, K. (1997). Changes of ethylene evolution, ACC content, ethylene forming enzyme activity and respiration in fruits of highbush blueberry. J. Japan Soc. Hort. Sci. 66, 23–27. doi:10.2503/jjshs.66.23

Tsuchiya, M., Satoh, S., and Iwai, H. (2015). Distribution of XTH, expansin, and secondary-wall-related CesA in floral and fruit abscission zones during fruit development in tomato (*Solanum lycopersicum*). Front. Plant Sci. 6, 323. doi:10.3389/fpls.2015.00323

[Dataset] USDA Economic Research Service (2020). Fruit and vegetable prices. Accessed: 2020-02-10

Vashisth, T., NeSmith, D. S., and Malladi, A. (2015). Anatomical and gene expression analyses of two blueberry genotypes displaying differential fruit detachment. J. Amer. Soc. Hort. Sci. 140, 620–626. doi:10.21273/JASHS.140.6.620

Vicente, A. R., Ortugno, C., Rosli, H., Powell, A. L., Greve, L. C., and Labavitch, J. M. (2007). Temporal sequence of cell wall disassembly events in developing fruits. 2. Analysis of blueberry (*Vaccinium* species). J. Agric. Food Chem. 55, 4125–4130. doi:10.1021/jf063548j

Wang, X., Liu, D., Li, A., Sun, X., Zhang, R., Wu, L., et al. (2013). Transcriptome analysis of tomato flower pedicel tissues reveals abscission zone-specific modulation of key meristem activity genes. PLoS ONE 8. doi:10.1371/journal.pone.0055238

Xin, Z. and Chen, J. (2012). A high throughput DNA extraction method with high yield and quality. Plant Methods 8, 26. doi:10.1186/1746-4811-8-26

Yu, P., Li, C., Takeda, F., and Krewer, G. (2014). Visual bruise assessment and analysis of mechanical impact measurement in southern highbush blueberries. Appl. Eng. Agric. 30, 29–37. doi:10.13031/aea.30.10224

Zhu, Y., Zheng, P., Varanasi, V., Shin, S., Main, D., Curry, E., et al. (2012). Multiple plant hormones and cell wall metabolism regulate apple fruit maturation patterns and texture attributes. Tree Genet Genome 8, 1389–1406. doi:10.1007/s11295-012-0526-3

